# Optimizing miRNA-Based Breast Cancer Subtyping with AHALA: A Multi-Stage Classification Approach

**DOI:** 10.1101/2025.04.03.646979

**Authors:** Mohammed Qaraad, Eric Rahrmann, David Guinovart

## Abstract

This paper presents the Adaptive Hill Climbing Artificial Lemming Algorithm (AHALA), a novel optimization method designed to improve miRNA-based breast cancer subtyping and classification. By integrating Hill Climbing into the Artificial Lemming Algorithm, AHALA balances global exploration and local exploitation through adaptive boundary handling and dynamic step-size decay. Applied to breast cancer miRNA expression data, AHALA excelled in multi-classification tasks, achieving an accuracy of 95.74%, precision of 95.98%, recall of 95.74%, F1 score of 95.74%, and AUC of 0.9682. These results highlight its ability to distinguish between breast cancer subtypes, including Luminal A, Luminal B, HER2-enriched, and Basal-like. Feature selection powered by AHALA identified key miRNAs—hsa-miR-190b, hsa-miR-429, hsa-miR-505-3p, hsa-miR-3614-5p, and hsa-miR-935—linked to subtype differentiation. Additionally, AHALA optimized neural network hyperparameters, including hidden layer size, learning rate, and batch size, enhancing classification performance. These findings demonstrate AHALA’s potential as a robust tool for miRNA classification, biomarker discovery, and machine learning optimization in breast cancer research, contributing to improved diagnostics and treatment strategies.

## 1. Introduction

Breast cancer is a highly heterogeneous disease that is commonly classified into four molecular subtypes based on the expression levels of hormone receptors and human epidermal growth factor receptor 2 (HER2): Luminal A, Luminal B, HER2-enriched, and Basal-like subtypes [1]. Each subtype exhibits distinct clinical outcomes, prognoses, and therapeutic responses [2]. Luminal A tumors, which are generally hormone receptor-positive and HER2-negative, tend to grow more slowly and are associated with a favorable prognosis, making them highly responsive to hormone therapies such as tamoxifen or aromatase inhibitors [3]. Luminal B tumors, while also hormone receptor-positive, demonstrate higher proliferation rates, often necessitating more aggressive treatment regimens, including chemotherapy in addition to hormone therapy [4]. HER2-enriched breast cancers, characterized by the overexpression of the HER2 protein and typically lacking hormone receptors, are more aggressive but respond well to HER2-targeted treatments such as trastuzumab, which has significantly improved patient outcomes [5]. On the other hand, Basal-like breast cancers, which show gene expression patterns resembling basal epithelial cells and lack both hormone receptors and HER2 expression, represent a highly aggressive and heterogeneous subtype. Chemotherapy remains the primary treatment option for this group due to the absence of targeted therapies [6]. Identifying and understanding these molecular subtypes is critical for developing personalized treatment strategies, and ongoing research aims to refine these classifications further and develop novel therapeutic approaches for improved patient outcomes across all breast cancer subtypes. The roles of microRNAs (miRNAs) in breast cancer biology are both significant and intricate. miRNAs are short non-coding RNAs that regulate gene expression and are implicated in various biological processes, including cancer development and progression.

Several studies have indicated that miRNAs hold great promise as potential biomarkers for cancer diagnosis, prognosis, and therapy [7–9]. Identifying these miRNA biomarkers is crucial due to their role in regulating gene expression, with their dysregulation being a common feature in many cancers. In cancerous cells, miRNAs can be overexpressed, acting like oncogenes by down-regulating tumor suppressor genes, which ordinarily work to prevent uncontrolled cell growth. Conversely, miRNAs that normally act as tumor suppressors may be under-expressed, leading to the upregulation of oncogenes, further driving cancer progression [10]. This dual role of miRNAs in both up-regulation and down-regulation makes them valuable as biomarkers for cancer. However, identifying specific miRNAs as biomarkers is complex, primarily due to the heterogeneous nature of cancer and the high dimensionality and noise present in miRNA expression datasets [11,12].

Numerous studies have focused on identifying miRNAs associated with breast cancer using a variety of datasets and methodological approaches, particularly in terms of feature selection. Common methods include Information Gain, Chi-Squared, and LASSO, combined with classification models like Random Forest and Support Vector Machines [13]. Despite the progress made, many of these studies have limitations, as they often focus only on miRNAs discovered through wet lab experiments, potentially overlooking recently identified miRNAs that could be effective biomarkers. Additionally, various ensemble feature selection techniques have been explored, including Stochastic Gradient Descent (SGD), Support Vector Machine classifier (SVC), gradient boosting, random forest, logistic regression, passive-aggressive classifier, ridge classifier, and bagging [14].

Other methods involve using ANOVA, Mutual Information, Extra Trees Classifier, Logistic Regression (LGR) [15], genetic algorithms [16], and LASSO [17]. However, much of this work is restricted to binary classification, such as distinguishing Triple-Negative Breast Cancer (TNBC) from non-TNBC [14], breast cancer from normal tissue [15,16], or breast cancer from other cancer types [17].

Moreover, some identified miRNAs, such as those reported in Ref. [15], have not undergone validation. Without enrichment analysis, there is a higher risk of false positives—miRNAs that may be statistically significant but lack biological relevance to breast cancer. Enrichment analysis is essential for filtering out such false discoveries, ensuring that the identified miRNAs are biologically meaningful and relevant. Among existing studies, only a few [18,19] have focused on the multi-class classification of breast cancer subtypes. In Ref. c, miRNAs were first filtered using XGBoost and Random Forest, and then the mutual information between molecules was calculated and used as a threshold in multilayer network analysis. However, the method to determine this threshold was overly complex, and no enrichment analysis was conducted. Similarly, the feature selection method in Ref. [18] was intricate, involving eight mutual information-based algorithms, such as MIM, mRMR, CMIM, JMI, DISR, ICAP, CIFE, and CONDRED. The feature subsets from each algorithm were carefully combined and used with the Random Forest classifier, resulting in a complicated but effective selection process.

In recent years, various methodologies have been applied to miRNA-based breast cancer classification, including traditional feature selection techniques like Information Gain and Chi-Squared [13], as well as more advanced machine learning algorithms such as Random Forests and Support Vector Machines [20]. While these methods have achieved notable successes, they often suffer from limitations in terms of scalability, robustness, and the ability to fully explore the complex miRNA interactions that define breast cancer subtypes. Furthermore, most existing studies focus on binary classifications, neglecting the multi-class nature of breast cancer subtyping [21]. To address these challenges, we propose a refined methodology to improve feature selection and breast cancer subtyping. Instead of employing association rule mining, we begin by removing low-variance genes from the dataset. The rationale behind this step is that genes with low variance across samples are less likely to be informative for classification, as they do not exhibit significant changes in expression that could distinguish between different cancer subtypes. Eliminating such genes reduces the dimensionality of the data and improves the overall efficiency of subsequent analyses. Following this, we apply differential gene expression (DGE)[22] analysis to identify genes that are significantly differentially expressed between breast cancer subtypes. DGE allows us to pinpoint biomarkers that may play crucial roles in distinguishing between subtypes based on their expression profiles, thereby enhancing the biological relevance of our feature set. This step is critical as it helps focus the classification task on the most biologically significant genes, rather than including a broad range of potentially irrelevant features. Once the key genes are selected, we introduce a novel optimization algorithm, the Adaptive Hill Climbing Artificial Lemming Algorithm (AHALA), to fine-tune a deep neural network designed for breast cancer subtype classification. AHALA improves both the exploration of the search space and the exploitation of optimal solutions, ensuring that the deep neural network is well-optimized to classify subtypes such as Luminal A, Luminal B, HER2-enriched, and Basal-like with high accuracy. The deep neural network is then trained on the differentially expressed genes, leveraging AHALA’s optimization capabilities to achieve superior classification performance.

To interpret the results of this classification task, we extract the most important biomarkers that contribute to the accurate prediction of breast cancer subtypes. This step enables us to identify critical miRNAs or genes that are key indicators of specific subtypes, offering valuable insights into the molecular mechanisms underlying breast cancer and potential targets for therapy.

In this research, we introduce an innovative approach for the identification of prospective microRNA (miRNA) biomarkers associated with the four primary molecular subtypes of mammary carcinoma: Luminal A, Luminal B, Human Epidermal growth factor Receptor 2 (HER2)-enriched, and Basal-like. The study utilizes a publicly accessible dataset comprising miRNA expression profiles and corresponding clinical information, as curated by previous researchers [19]. This comprehensive dataset incorporates next-generation sequencing (NGS)-derived miRNA expression values alongside pertinent clinical data, all of which have been sourced from The Cancer Genome Atlas (TCGA) repository.

The main contributions of this work include:

- Development of AHALA: A novel hybrid optimization algorithm balancing global exploration and local exploitation for improved feature selection and classification.
- AHALA application to breast cancer subtyping: Fine-tuning a deep neural network for accurate classification of breast cancer subtypes using differentially expressed genes.
- Key biomarker identification: Utilizing AHALA to pinpoint critical miRNAs and genes associated with breast cancer subtypes, offering potential diagnostic and therapeutic targets.
- Validation on CEC2021 benchmark: Demonstrating AHALA’s versatility and effectiveness as a general-purpose optimization technique for diverse, complex challenges.

Through extensive experimentation on breast cancer datasets, AHALA demonstrates significant improvements in classification accuracy, precision, recall, and F1-score, highlighting its potential as a powerful tool in breast cancer subtyping and biomarker discovery. The remainder of this paper is organized as follows: Section 2 details the methodology and data used in our study, Section 3 presents the results, Section 4 discusses the biological significance of our findings, and Section 5 concludes with future directions for research.

## 2. Methodologies

In this section, we outline the methodologies employed, beginning with an overview of the Artificial Lemming Algorithm (ALA) process, followed by a detailed description of the proposed Adaptive Hill Climbing Artificial Lemming Algorithm (AHALA) and its enhancements for optimization tasks.

### 2.1 Artificial lemming algorithm (ALA)

Arctic ecosystems are home to a remarkable micromammal known as the lemming, a member of the Cricetidae family. These small rodents, typically measuring around 10 centimeters in length, are primarily found in tundra, forest-tundra, and mountain-tundra habitats [23]. Lemmings exhibit a distinctive morphology, characterized by a rotund body shape and pelage that varies in coloration depending on the species, ranging from grey to yellow and brown hues. One of the lemming’s most notable anatomical features is a specialized flattened claw on the first digit of their anterior appendages, which aids in excavating snow-covered terrain. Their dietary habits are predominantly herbivorous, with a preference for graminoids and bryophytes. However, they display dietary flexibility when necessary, consuming foliage, fruits, bulbous structures, roots, and even lichens found atop the snow layer [24]. Within the Arctic food web, lemmings occupy a precarious position, facing constant predation pressure from various carnivores, including Bubo scandiacus (snowy owls), mustelids (weasels), and Ursus maritimus (polar bears). Despite these challenges, lemmings have evolved an extraordinary reproductive capacity. They can produce up to eight litters annually, with each litter potentially yielding up to a dozen offspring. Furthermore, their sexual maturation occurs rapidly, with individuals attaining reproductive capability within a mere 20 to 30 days post-partum [25]. To support their prodigious reproductive output, lemmings have developed the ability to consume quantities of food equivalent to twice their body mass in a single feeding session. Their annual behavioral patterns include springtime movements to higher elevations, where they inhabit mountainous areas and forested regions. During this period, they engage in continuous reproduction before returning to their tundra habitats in autumn [26]. Lemmings rely heavily on their acute olfactory and auditory senses to locate sustenance both above and below ground. When confronted with predators, they employ a diverse repertoire of evasive strategies. As skilled fossorial animals, lemmings construct intricate subterranean networks of tunnels and chambers, which serve dual purposes as secure dwellings and food storage facilities [27]. The ALA algorithm’s mathematical model is presented as follows:

#### 2.1.1 Initialization

The Artificial Lemming Algorithm (ALA) is a population-based optimization method that begins by initializing the positions of search agents. These initial candidate solutions form a matrix ***Z***, with ***N*** rows (population size) and ***Dim*** columns (number of dimensions), constrained by the problem’s upper and lower bounds, as outlined in Equation (1). The best solution found during each iteration is considered the optimal or near-optimal result at that stage. Decision variables ***z***_***i***,***j***_ for each dimension are calculated using Equation (2).

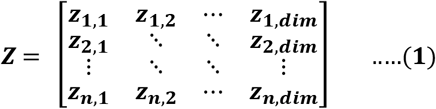

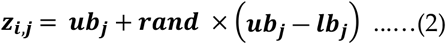

In the Artificial Lemming Algorithm (ALA), the random value ***rand*** falls within the range [0, 1]. The variables ***lb***_***j***_ and ***ub***_***j***_ represent the lower and upper bounds, respectively, for the ***j***^***th***^ dimension. These values are used to calculate the decision variables ***z***_***i***,***j***_, which define the positions of the search agents within the specified limits of each dimension.

#### 2.1.2 Long-Distance Migration (Exploration)

In the first behavioral phase of the Artificial Lemming Algorithm (ALA), lemmings randomly perform long-distance migrations when food becomes scarce due to overpopulation. During this exploration phase, lemmings move through the search space, guided by their current position and the positions of random individuals within the population. This migration aims to discover regions with richer resources and improved living conditions. Notably, the direction and distance of the migration vary depending on ecological factors, making the movement dynamic. This behavior is mathematically modeled by the following equation:

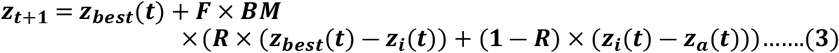

In this model, ***z***_***t***+1_ represents the updated position of the ***i***^***th***^search agent at iteration(t+1), while ***z***_***best***_**(*t*)** indicates the best solution found so far. The variable ***F***, introduced in Equation (5), serves as a flag that alters the search direction to prevent the algorithm from getting stuck in local optima, thus ensuring more comprehensive exploration of the search space. ***BM*** refers to a vector of random numbers simulating Brownian motion, which introduces dynamic and uniform step sizes, helping the search agents explore potential regions more effectively. The step size for Brownian motion is determined by the probability density function of a normal distribution with a mean of 0 and variance of 1, as defined in Equation (4). Additionally, ***R*** is a vector of size 1 × ***Dim*** whose elements are random values uniformly distributed within the range [-1, 1], generated using Equation (6).

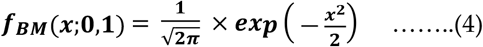

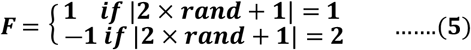

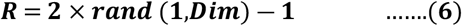

#### 2.1.3 Digging Holes (Exploration)

The second behavior in the Artificial Lemming Algorithm (ALA) models lemmings digging burrows in their environment, forming intricate tunnel networks for protection and food storage. Lemmings create new burrows randomly, guided by their current position and the locations of other random individuals in the population. This behavior enhances their ability to evade predators and search for food more efficiently. The digging process is mathematically modeled by Equation(7).

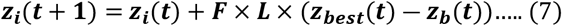

In this context, ***L*** represents a random number related to the current iteration count, influencing the interactions between lemming individuals during the burrow-digging process. ***z***_***b***_ denotes a randomly selected individual from the population, with bbb being a randomly chosen integer index between 1 and ***N*** (population size). These variables ***L*** and ***z***_***b***_ model the cooperative behavior of lemmings as they dig new burrows. The value of ***L*** is calculated according to the following equation:

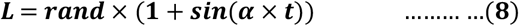

***α*** represents a small fixed value equal to 0.5.

#### 2.1.4 Foraging for Food (Exploitation)

In the third behavior, lemmings engage in extensive, random movements within their burrow networks, utilizing their acute senses of smell and hearing to detect food. They typically confine their foraging efforts to a relatively small area based on the availability and abundance of food. To maximize their intake, lemmings wander randomly within these foraging zones. This stage of behavior is modeled using a spiral wrapping mechanism, which simulates the lemmings’ movement pattern as follows:

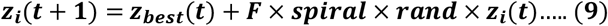

In this model, the term “spiral” represents the spiral-shaped trajectory of the lemmings’ random search during foraging. This pattern simulates their movement as they search for food within a limited area. The spiral movement is mathematically described by Equations (10) and (11), which govern the shape and behavior of the foraging process.

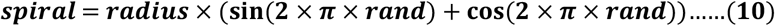

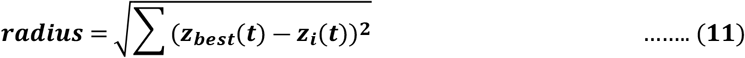

In this context, “radius” refers to the radius of the lemmings’ foraging area, defined as the Euclidean distance between their current position and the optimal solution. This radius helps determine the extent of the search area during foraging, influencing the lemmings’ ability to locate food sources effectively.

#### 2.1.5 Evading Natural Predators (Exploitation)

In the final stage, the model emphasizes the avoidance and protective behaviors of lemmings in the face of danger. The burrow acts as a safe refuge for these animals. Upon detecting a predator, lemmings utilize their remarkable running speed to retreat quickly to their burrow. Additionally, they employ deceptive maneuvers to evade capture during pursuit. This behavior is mathematically represented by Equation (12).

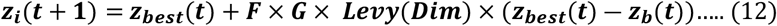

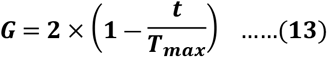

In this context, ***G*** denotes the escape coefficient of lemmings, which quantifies their ability to flee and diminishes as the iteration count increases, as indicated in Equation (13). ***T***_***max***_ represents the maximum number of iterations for the algorithm. The Lévy flight function, denoted as ***Levy*(** · ), is utilized to model the deceptive maneuvers lemmings execute while escaping predators. This function is expressed as follows, where *u* and *v* are random values within the interval [0, 1], and ***β*** is a constant set to 1.5.

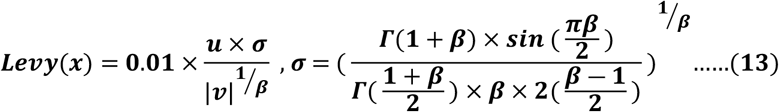

In the Artificial Lemming Algorithm (ALA), the four search strategies are directly linked to the energy levels of the lemmings. During the initial phases, lemmings primarily focus on exploration to identify potential areas. As the search progresses, they shift towards local exploitation to refine their solutions. To achieve a balanced approach between exploration and exploitation, an energy factor is introduced, which decreases over successive iterations. When lemmings possess adequate energy, they actively engage in migration or burrow digging; conversely, when energy is low, they prioritize foraging and evading predators. The formula for calculating the energy factor is as follows:

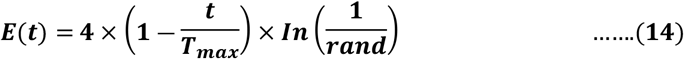

***t*** represents the current iteration, ***T***_***max***_denotes to the maxmum number of iterations, and ***rand*** represents random number in range [0,1].

### 2.2 Proposed Adaptive Hill Climbing Artificial Lemming Algorithm AHALA

The Artificial Lemming Algorithm (ALA) is a bionic meta-heuristic optimization technique inspired by the behavior of lemmings. While ALA has shown promise in solving complex optimization problems, there is potential for improvement in its convergence speed and solution quality. Meta-heuristic algorithms often face challenges balancing exploration (searching global space) and exploitation (refining local solutions). ALA’s strength lies primarily in its exploration capabilities, but it may benefit from enhanced local search mechanisms to improve its exploitation phase.

#### 2.2.2 Insight of Enhancement

The proposed enhancement introduces a local search component, specifically a Hill Climbing algorithm, to complement ALA’s global search strategy. This hybridization aims to leverage the strengths of both approaches:

1. ALA’s global search capability helps to explore the solution space broadly and avoid getting trapped in local optima.
2. The Hill Climbing algorithm provides a mechanism for fine-tuning promising solutions, potentially accelerating convergence and improving the quality of final solutions.

By periodically applying local search to the best solution found by ALA, we create opportunities for incremental improvements that might be overlooked by the main algorithm’s larger steps.

#### 2.2.2 Mathematical Description

Let ***f***: ℝ^***n***^ → ℝ be the objective function to be minimized, where ***n*** is the dimension of the problem space. The enhanced AHA (Adaptive Hill Climbing Artificial Lemming Algorithm) can be described as follows:

Initialization: Initialize a population of ***N*** solutions: ***X* = {*x***_1_, ***x***_**2**_, **…, *x***_***n***_**}**, where each ***x***_***n***_ **∈** ℝ^***n***^.

Main Iteration Loop: For each iteration ***t* =** 1,**2**,**…**,***T***

a. Apply the standard AHA update rules to generate new candidate solutions.
b. Evaluate the fitness of the new solutions: ***f*(*x***_***i***_) for each ***x***_***i***_ **∈ *X***.
c. Update the best-known solution ***x***_***best***_
d. If ***t*** mod ***k* = 0** (where ***k***is a predefined frequency), apply Hill Climbing to ***x***_best_.

Hill Climbing Procedure: For ***j* =** 1,**2**,**…**,***M***(where ***M***is the maximum number of Hill Climbing iterations):

a. Generate a perturbation vector ***δ***_***j***_ **∈** ℝ^***n***^, where each component is drawn from a uniform distribution: ***δ***_***j***_ **∼ *U* (** − 1,1)
b. Create a new candidate solution:

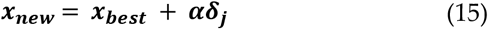

where ***α*** is the step size.
c. If ***f*(*x***_***new***_**) *< f*(*x***_***best***_), update ***x***_***best***_ **= *x***_***new***_.
d. Otherwise, reduce the step size:

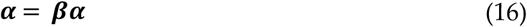

where **0 *< β <*** 1is a decay factor. ***k*** represents the Frequency of applying the Hill Climbing procedure, such as every 10 iterations of the AHA algorithm. ***M*** denotes the Maximum number of Hill Climbing iterations, ***α*** represents Initial step size for Hill Climbing, and ***β*** refer to the Step size decay factor. The choice of ***k*** plays a critical role in balancing exploration and exploitation. If ***k*** is too small (i.e., Hill Climbing is applied too frequently), the algorithm may focus too heavily on local exploitation, potentially neglecting the broader search space and leading to premature convergence. On the other hand, if ***k*** is too large (i.e., Hill Climbing is applied infrequently), the algorithm might miss opportunities to refine the best solution found so far, slowing down convergence.

Similarly, the initial step size ***α*** and the decay factor ***β*** are crucial in controlling the granularity of local search during Hill Climbing. A larger ***α*** allows for more significant perturbations, encouraging broader exploration within the neighborhood of the current best solution. However, if ***α*** is too large, the algorithm might overshoot optimal solutions. The decay factor ***β*** gradually reduces the step size over time, allowing the search to focus more finely on local optima as the algorithm progresses. This decaying step size ensures that, as the search continues, the adjustments become more precise, improving the likelihood of finding a near-optimal solution. Together, these parameters provide flexibility, enabling the algorithm to fine-tune the balance between global search and local refinement, which is essential for efficiently navigating complex, high-dimensional search spaces.

#### 2.2.3 Complexity Analysis

The computational complexity of the Adaptive Hill Climbing Artificial Lemming Algorithm (AHALA) can be understood by analyzing both the standard AHA operations and the additional computational load introduced by the Hill Climbing mechanism.

##### Standard AHA Complexity

The standard AHA optimization process involves maintaining a population of ***N*** candidate solutions, each residing in an ***n*** − ***dimensional*** space. The algorithm iterates over a total of ***T*** iterations, and the computational cost of evaluating the objective function ***f*** for each candidate, the solution is denoted by ***O*(*E*)**, where ***E*** represents the cost of a single fitness evaluation. Thus, the total time complexity for the standard AHA algorithm can be expressed as: ***O* (*NTnE*)** This term captures the cost of evaluating all candidate solutions over ***T*** iterations across ***n*** − ***dimensional*** problem space.

##### Hill Climbing Enhancement Complexity

The integration of the Hill Climbing mechanism into AHA introduces additional computational costs. Hill Climbing is applied periodically, once every ***k*** iterations, to refine the best-known solution ***x***_***best***_. For each application of Hill Climbing, a maximum of ***M*** iterations are performed, where each iteration requires the generation of a perturbation vector in the ***n*** − ***dimensional*** space and a fitness evaluation. The computational complexity of the Hill Climbing phase per application is thus ***O*(*MnE*)**, and since Hill Climbing is applied 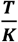 times during the entire optimization process, the total additional complexity is given by:

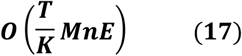

##### Overall Complexity

Combining the time complexity of the standard AHA and the additional cost introduced by Hill Climbing, the overall time complexity of the AHALA can be expressed as:

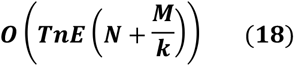

This expression indicates that the computational load grows linearly with the number of iterations ***T***, the dimensionality of the problem ***n***, and the cost of evaluating the fitness function ***E***. The terms ***N*** (population size) and 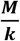 (related to the frequency and intensity of Hill Climbing) dictate the relative contributions of global exploration and local exploitation to the overall complexity.

##### Space Complexity

The space complexity of AHALA remains largely unchanged from the standard AHA algorithm. Since the algorithm only requires storage for the population of candidate solutions and a constant number of variables used for Hill Climbing, the overall space complexity is: ***O*(*Nn*)** This space complexity is proportional to the number of candidate solutions ***N*** and the dimensionality of the search space ***n***, with no significant additional storage required for the hill-climbing process. The proposed enhancement introduces a trade-off between increased computational cost and potential improvements in convergence speed and solution quality. The additional complexity of 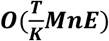 is generally acceptable if ***M*** and 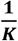 are kept reasonably small. By carefully tuning these parameters, the computational overhead remains manageable while still achieving significant gains in performance. The effectiveness of this enhancement may vary depending on the nature of the optimization problem. It is particularly beneficial for problems with the following characteristics:

- Rugged fitness landscapes with many local optima
- Problems where small, local improvements can significantly impact the overall solution quality
- Scenarios where the computational budget allows for additional local search steps

The parameters ***k, M***,***α*** and ***β*** provide flexibility in balancing the global exploration of ALA with the local exploitation of Hill Climbing. These parameters can be tuned based on the specific problem characteristics and computational resources available. Future work could explore adaptive mechanisms for automatically adjusting these parameters during the optimization process, potentially leading to a more robust and efficient algorithm across a wider range of problem types. Algorithm 1 presents the pseudocode for the AHALA algorithm, while Figure 1 provides a visual representation of the algorithm’s procedural flow.

**Fig. 1.**
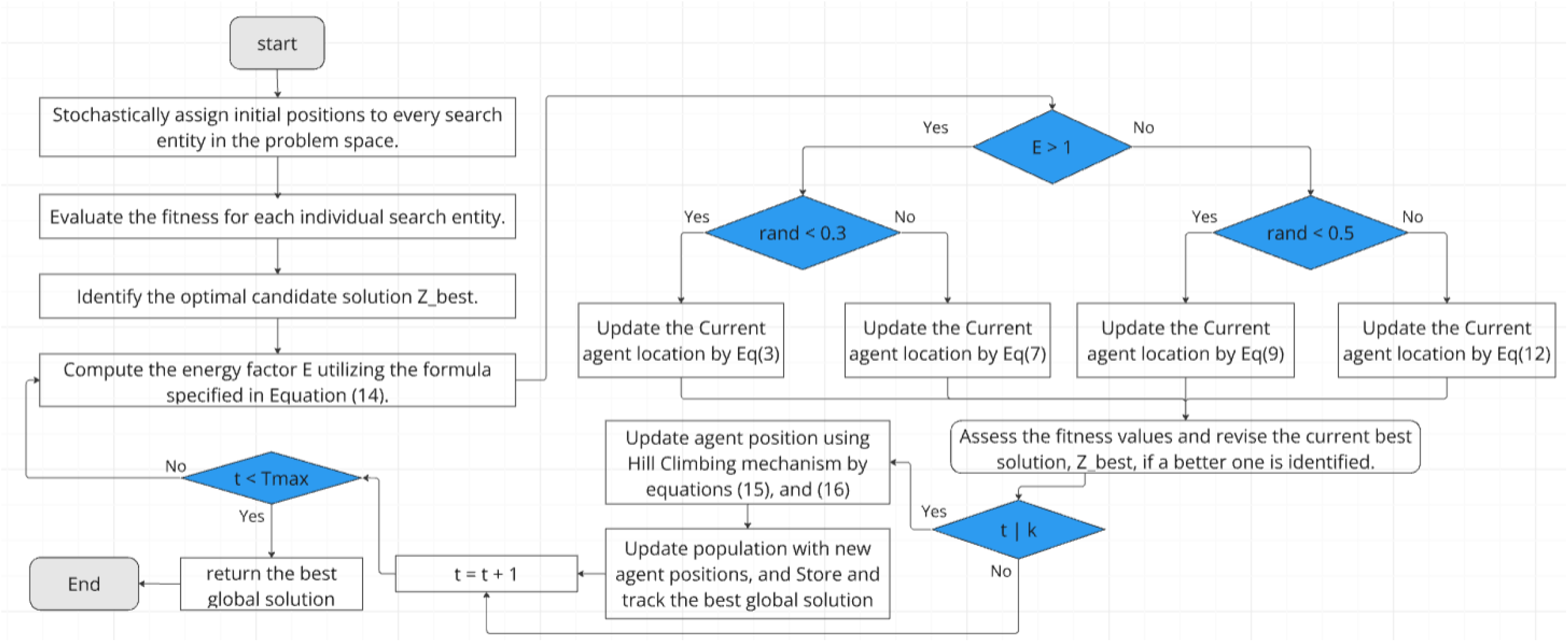
AHALA flowchart

###### Algorithm 1

The AHALA

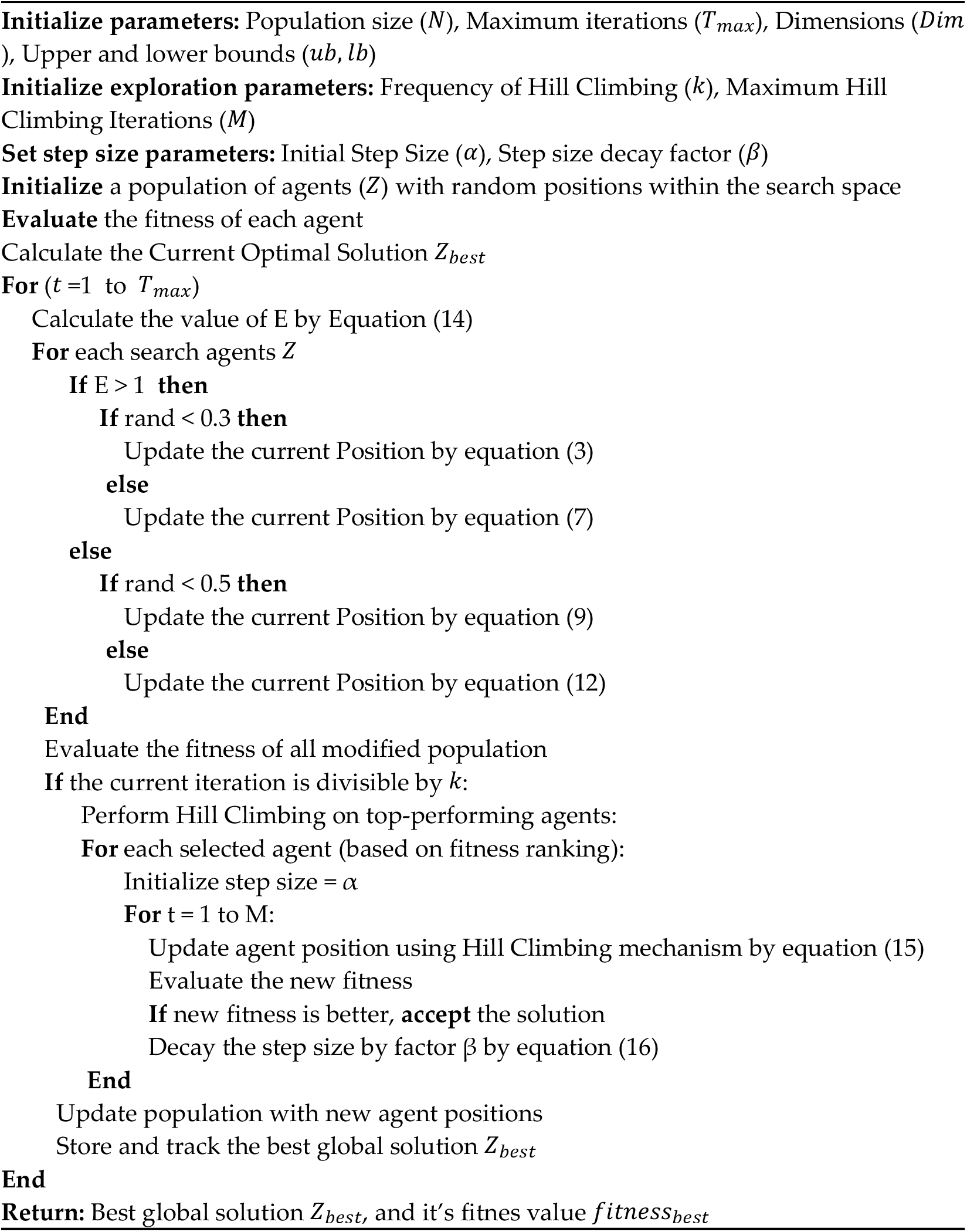

## 3. Experimental Validation and Performance Analysis

To evaluate the robustness and effectiveness of the proposed Adaptive Hybrid Artificial Lemming Algorithm (AHALA), we conducted comprehensive computational experiments utilizing the CEC2021 benchmark suite. This established benchmark framework provides a rigorous testing environment for assessing optimization algorithms across various complexity levels, offering challenging scenarios that effectively test both established and emerging metaheuristic approaches [28]. To establish a comprehensive performance baseline, we implemented comparative analyses against six contemporary metaheuristic algorithms developed between 2020 and 2024. The comparison group consisted of:

- The RIME algorithm[29].
- Q-learning embedded sine cosine algorithm (QleSCA) [30].
- Enhanced Tug of War Optimization(EnhancedTWO) [31].
- Random Walk Grey Wolf Optimizer (RWGWO) [32].
- Gradient-based optimizer(GBO) [33].
- Standard Artificial Lemming Algorithm (ALA).

For experimental consistency, we maintained uniform initial conditions across all algorithms. The population size (N) was standardized at 30 individuals, with a maximum evaluation threshold of 2500 iterations. Implementation parameters for each comparative algorithm were configured according to their respective original publications, as detailed in Table 1. The experimental framework utilized Python implementations executed on a high-performance computing environment running Rocky Linux 9.4 (Blue Onyx). The hardware configuration comprised an AMD EPYC 7713 64-core Processor supported by 1TiB RAM, ensuring consistent computational capabilities throughout the testing phase. To account for the stochastic nature of these algorithms, we executed 30 independent trials for each method. Performance metrics were calculated using multiple statistical indicators:

**Table 1:** Algorithm-Specific Parameter Configurations.

| Algorithm | parameters Specifications |
| --- | --- |
| RIME | Soft-rime(sr)=5.0; popSize=30; |
| EnhancedTWO | popSize=30; |
| GBO | popSize=30; |
| QleSCA | learning rate in Q-learning (alpha)=0.1;<br>(gama)=0.9; |
| RWGWO | popSize=30; |
| AHALA | max_iterations=10, step_size=0.01 |

- Mean fitness values (Avg)
- Standard deviation (std)
- Median performance (Med)
- Friedman rank statistics
- Wilcoxon signed-rank test results

### 3.1 Qualitative analysis

In this analysis, the performance of AHALA was evaluated on six CEC2021 benchmark functions (F1, F2, F3, F4, F9, and F10) as shown in Figure 2, each presenting unique optimization challenges. The analysis is based on six key performance indicators: function landscape, search history, diversity measurement, runtime, exploration vs. exploitation balance, and global best fitness.

**Fig 2.**
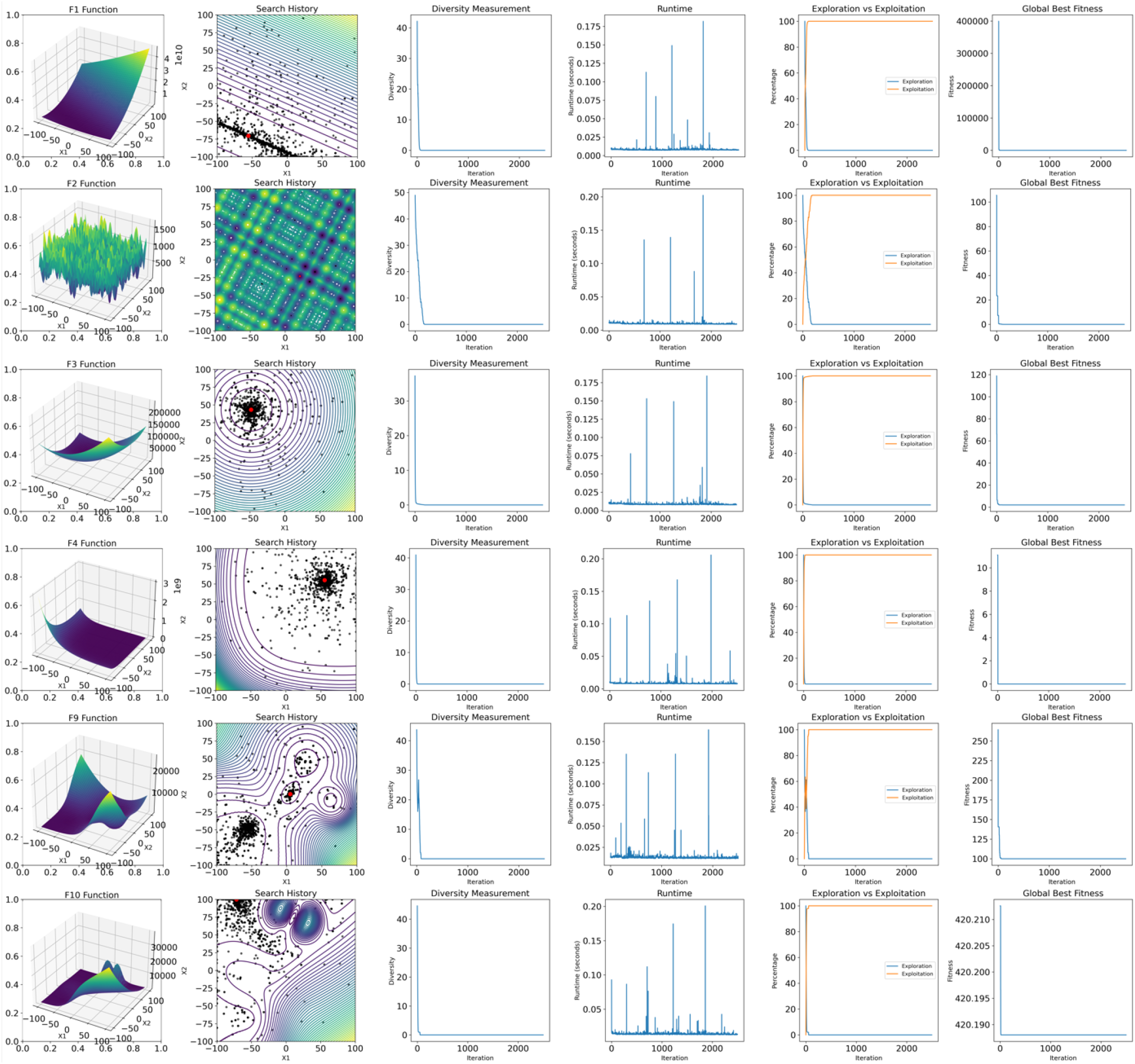
Qualitative analysis of AHALA on six CEC2021 benchmark functions, illustrating 3D surface plots, search history, diversity measurements, runtime behavior, exploration-exploitation balance, and global best fitness over iterations.

Function Landscape and Search History: The 3D surface plots and 2D contour plots reveal the diverse nature of the benchmark functions. F1 and F4 appear to be unimodal, while F2, F3, F9, and F10 exhibit multimodal characteristics with varying degrees of complexity. The search history overlaid on the contour plots demonstrates AHALA’s ability to navigate these landscapes effectively. For instance, in F1 and F4, the algorithm converges quickly to the global optimum region, whereas in more complex functions like F9 and F10, it explores multiple promising areas before convergence.

Diversity Measurement: Across all functions, AHALA maintains a relatively high diversity in the initial stages, which gradually decreases as the search progresses. This trend is particularly pronounced in F1 and F4, indicating rapid convergence. In contrast, for multimodal functions like F2 and F10, the diversity reduction is more gradual, suggesting sustained exploration of the search space, which is a key benefit of its adaptive nature and hill-climbing exploitation mechanisms.

Runtime Performance: The runtime charts show sporadic spikes across all functions, which could indicate periodic intensive computations or adaptive mechanisms within AHALA. The frequency and magnitude of these spikes vary across functions, potentially reflecting the algorithm’s responsiveness to different landscape complexities. The relatively consistent runtime across iterations implies that the algorithm is not heavily burdened by significant fluctuations in computational cost, making it suitable for handling high-dimensional problems.

Exploration vs. Exploitation Balance: A common pattern observed across all functions is a high initial exploration phase followed by a transition to exploitation. The transition point and the rate of change vary depending on the function complexity. For simpler functions like F1, the transition occurs earlier, while for more complex functions like F10, the exploration phase is prolonged. This adaptive control is crucial in avoiding stagnation in local minima, which can be observed by the balance of exploration and exploitation percentages over iterations.

Global Best Fitness: The convergence behavior of AHALA differs significantly across functions. For F1 and F4, rapid initial improvement is followed by fine-tuning. In contrast, functions like F2 and F10 show more gradual improvement, with occasional significant jumps in fitness, indicative of escaping local optima. The convergence pattern indicates that AHALA is robust against varying problem complexities.

The Proposed Algorithm AHALA demonstrates adaptive behavior across diverse function landscapes. It effectively balances exploration and exploitation, showing rapid convergence for simpler functions while maintaining the ability to navigate complex, multimodal spaces. The algorithm’s performance on these CEC2021 benchmarks suggests its potential versatility in handling a wide range of optimization problems. However, further quantitative analysis and comparison with other state-of-the-art algorithms would be necessary to fully assess its efficacy and efficiency.

### 3.2 Comparison with Recent Optimizers

The Discussion of Numerical Results highlights the comparative analysis of the proposed AHALA algorithm against six contemporary optimizers using the CEC2021 benchmark suite, with a focus on performance across various optimization scenarios (F1-F10). The findings reveal significant performance differences, as shown in Table 2, demonstrating AHALA’s strengths in global optimization and stability across different functions. In the context of global optimization, F1 Function, AHALA shows exceptional performance with an average value of 1.0280E+02, outperforming all competitors by several orders of magnitude. The closest competitor, ALA, achieves 1.4208E+03, while other algorithms, such as QleSCA, present much higher values (1.8437E+10). Statistical validation through the Wilcoxon signed-rank test supports this finding, consistently showing a p-value of 1.86E-09 when compared to all competitors, indicating that AHALA’s performance is statistically significant at the α=0.05 level, as shown in Table 3.

**Table 2:**
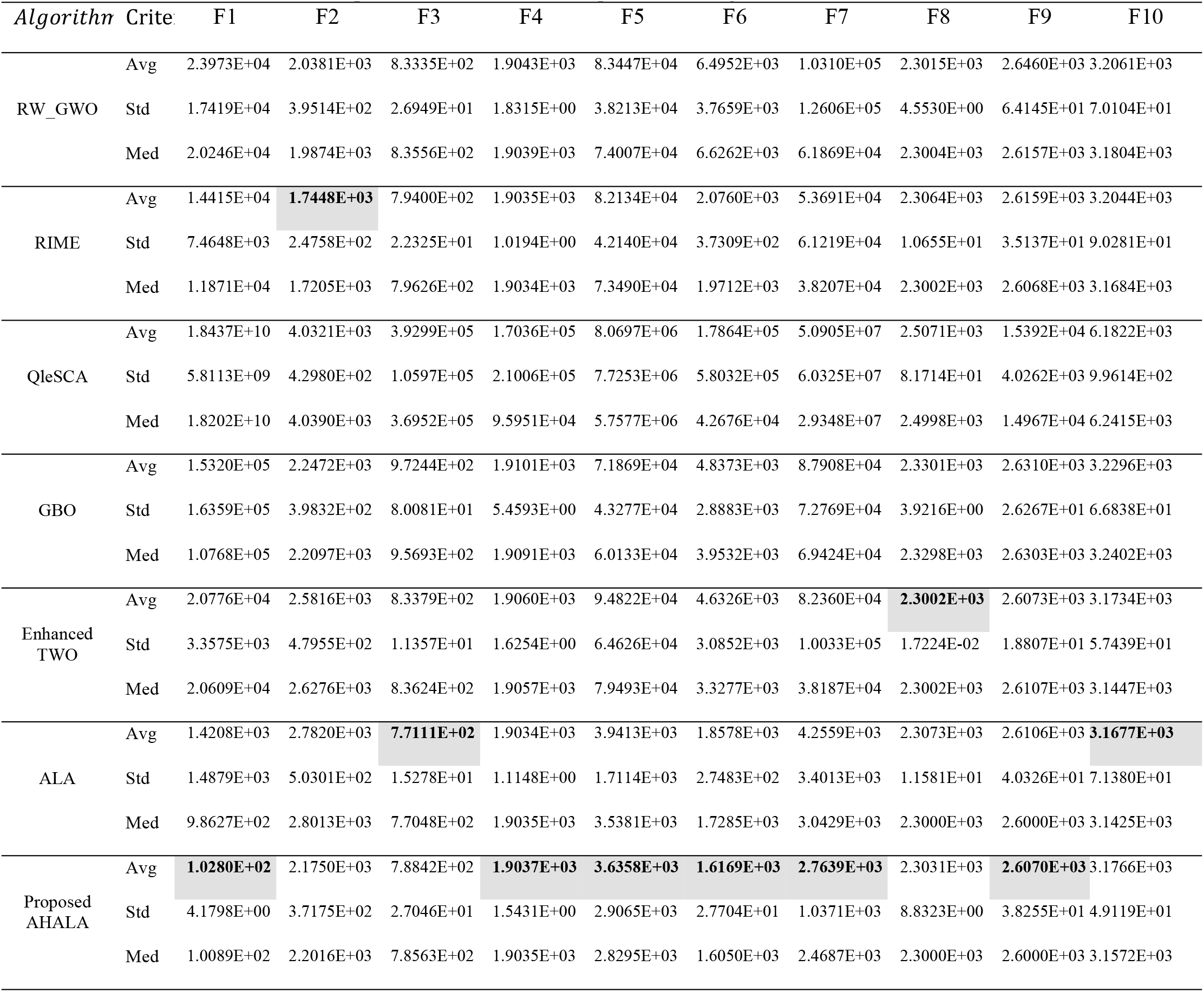
Comparison with some recent optimizers using CEC2021.

**Table 3:** Wilcoxon signed-rank test (p-value) results of AHALA and other algorithms on CEC2021.

| Function | Our Proposed AHALA Vs. |  |  |  |  |  |
| --- | --- | --- | --- | --- | --- | --- |
|  | ALA | Enhanced TWO | GBO | QleSCA | RIME | RWGWO |
| f1 | 1.86E-09 | 1.86E-09 | 1.86E-09 | 1.86E-09 | 1.86E-09 | 1.86E-09 |
| f2 | 8.86E-05 | 0.00202 | 0.626346 | 1.86E-09 | 9.90E-05 | 0.09195 |
| f3 | 0.01546 | 1.86E-08 | 1.86E-09 | 1.86E-09 | 0.489846 | 2.08E-05 |
| f4 | 0.427955 | 5.14E-06 | 2.05E-07 | 1.86E-09 | 0.776569 | 0.076721 |
| f5 | 0.02085 | 1.86E-09 | 1.86E-09 | 1.86E-09 | 1.86E-09 | 1.86E-09 |
| f6 | 0.000189 | 1.86E-09 | 1.86E-09 | 1.86E-09 | 1.86E-09 | 1.86E-09 |
| f7 | 0.004665 | 1.86E-09 | 1.86E-09 | 1.86E-09 | 1.86E-09 | 1.86E-09 |
| f8 | 0.100421 | 0.013663 | 3.73E-09 | 1.86E-09 | 0.005013 | 0.012834 |
| f9 | 1 | 0.000153 | 0.000137 | 1.86E-09 | 3.79E-06 | 3.79E-06 |

The stability and consistency of AHALA are also noteworthy, as demonstrated by its standard deviation metrics across functions. For instance, in F6, AHALA’s standard deviation (2.7704E+01) is substantially lower than that of other competitors, such as RW_GWO (3.7659E+03). Similarly, for F7, AHALA’s standard deviation (1.0371E+03) reflects greater stability when compared to RIME (6.1219E+04) and RW_GWO (1.2606E+05), underscoring the algorithm’s reliability in maintaining consistent results. In terms of comparative performance, AHALA showcases clear strengths across F1, F5, F6, and F7, where it significantly outperforms other algorithms. For example, in F1, AHALA’s median value (1.0089E+02) is drastically better than all other competitors. In contrast, functions like F4 and F9 show competitive but not superior performance, with p-values indicating statistical parity between AHALA, ALA (p=0.427955), and RIME (p=0.776569) for F4, and equivalent performance with ALA in F9 (p=1.000).

The statistical significance analysis further supports AHALA’s dominance in certain functions. For F1, the Wilcoxon signed-rank test reveals consistent superiority with p-values of 1.86E-09, as shown in Table 3) across all algorithm comparisons. In contrast, for F2, significance levels vary, ranging from p=1.86E-09 against QleSCA to p=0.626346 when compared to GBO. Additionally, AHALA shows statistically significant improvements in F5-F7 with strong p-values across all competitors. Several notable performance patterns emerge from this analysis. AHALA demonstrates consistent optimization efficiency, particularly in complex functions like F1, F5, and F6, where it consistently achieves lower average values. Furthermore, the algorithm strikes a balance between exploration and exploitation, a critical factor in ensuring efficient convergence. AHALA also exhibits convergence stability, with lower standard deviations across most functions, highlighting reliable performance. The fact that median values are consistently close to averages across different functions reflects AHALA’s robust convergence properties.

Despite these strengths, certain areas require further investigation. In particular, AHALA’s performance in F2 shows variability in significant levels, which could indicate areas for improvement. Additionally, while competitive, the algorithm does not demonstrate clear dominance in F4 and F9, warranting further exploration of its behavior in these functions.

Figure 3 presents the Friedman test results comparing AHALA with six other optimization algorithms on the CEC2021 benchmark. The Friedman test, a non-parametric statistical test, ranks the algorithms based on their performance across multiple functions. Lower ranks indicate better performance. The proposed AHALA algorithm achieved the second-best average rank (1.94), which demonstrates its competitive edge among the tested algorithms. The Adaptive Local Algorithm (ALA) obtained the best overall rank (2.26), suggesting it performed slightly better than AHALA in terms of consistency. Meanwhile, RIME was ranked third with a Friedman rank of 3.2, followed by EnhancedTWO and RWGWO with ranks of 4.2 and 4.21, respectively. The Gradient-Based Optimizer (GBO) and QleSCA performed less favorably, with GBO receiving a rank of 5.04 and QleSCA the worst rank of 7. These rankings reflect the overall robustness of AHALA across various test functions, positioning it as a highly competitive algorithm among the recent optimization methods.

**Fig 3:**
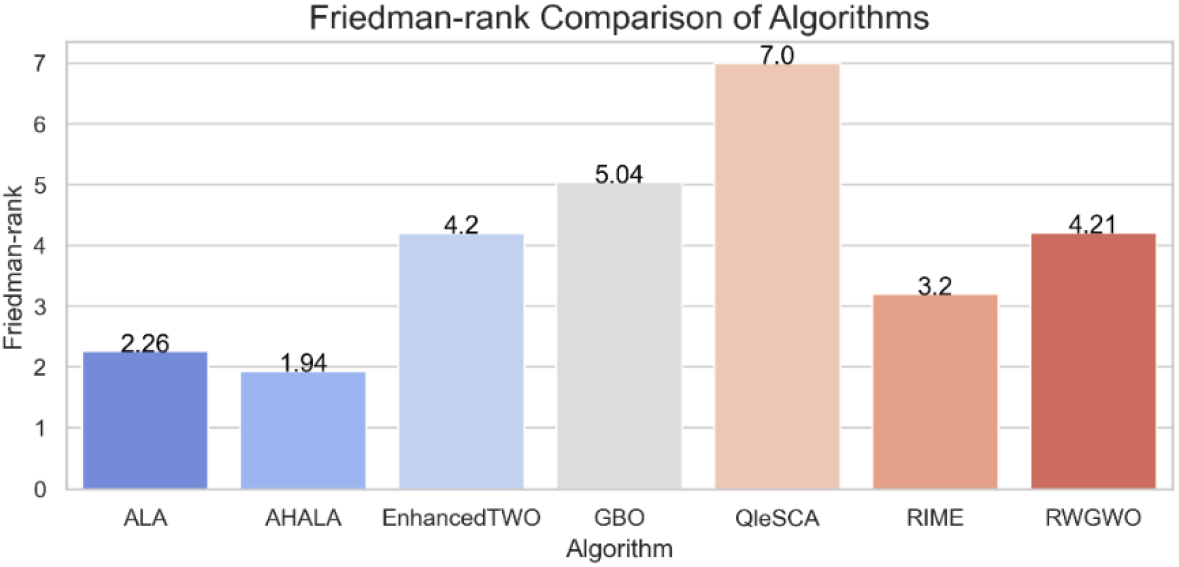
Friedman test results of AHALA and other algorithms on CEC2021

### 3.3 Convergence Analysis

In this subsection, we analyze the convergence behavior of the proposed AHALA algorithm in comparison to the basic ALA. Figure 4 illustrates the convergence curves of AHALA and ALA across multiple objective functions from the CEC2021 benchmark. The convergence analysis aims to evaluate the ability of the algorithms to minimize the fitness function over increasing iterations. For objective functions f1, f2, f5, and f7, AHALA demonstrates faster convergence, consistently achieving lower fitness values than ALA, particularly in the later stages of optimization. This indicates AHALA’s superior exploitation capability and its ability to effectively refine solutions as the iterations progress. Notably, for f2 and f5, the gap between AHALA and ALA becomes more pronounced during the final iterations, reflecting AHALA’s enhanced local search mechanism.

**Fig 4.**
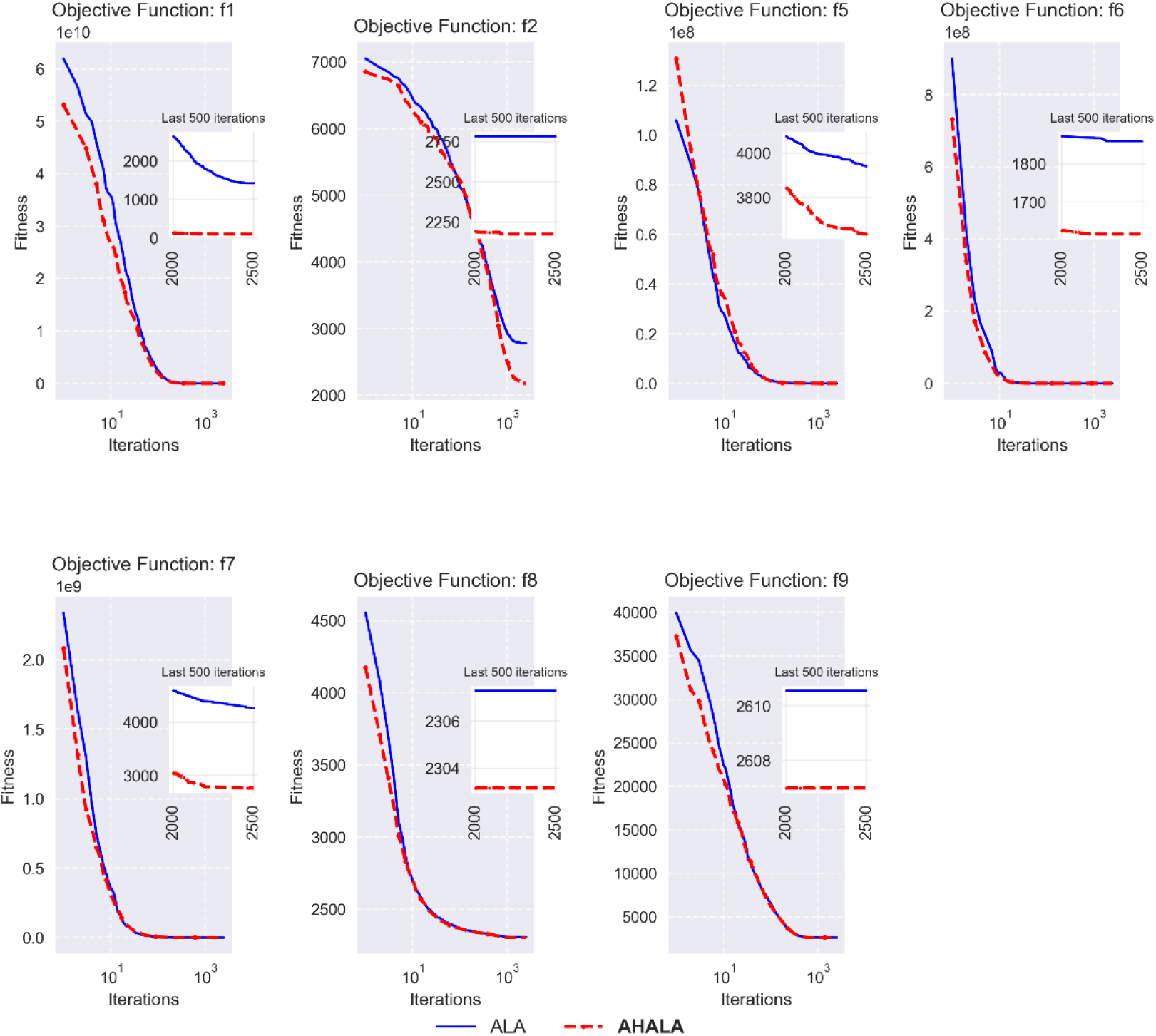
Convergence behavior of AHALA Vs basic ALA on some of the CEC2021 test suites

**Fig 4a.**
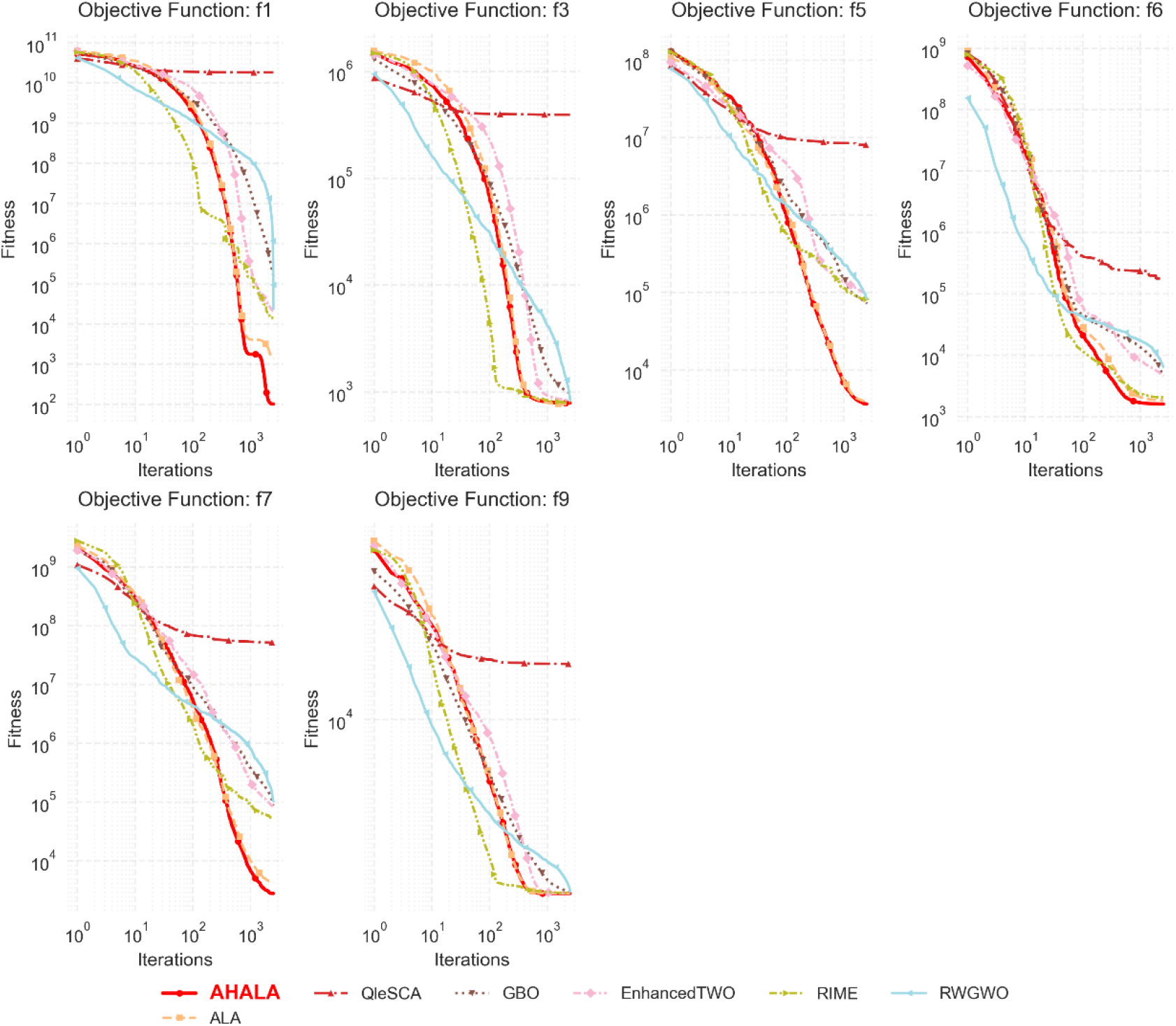
Convergence behavior of AHALA Vs other algorithms on some of the CEC2021 test suites

In contrast, for functions f6 and f9, AHALA and ALA exhibit similar convergence patterns, with both algorithms achieving comparable fitness values. This suggests that both algorithms are well-suited for optimizing these particular functions, although AHALA retains a slight edge in maintaining a more stable performance towards the end of the search process. The insets within each plot provide further insight into the final 500 iterations, where AHALA maintains a more consistent and lower fitness value across several functions, demonstrating its advantage in fine-tuning solutions during the final optimization phase. The reduced oscillations in the AHALA curve, as seen in functions like f7 and f8, further support its robust convergence behavior. Overall, AHALA outperforms ALA in most of the tested functions, particularly in terms of convergence speed and precision, making it a more effective algorithm for tackling high-dimensional optimization problems.

On the other hand, the convergence characteristics of the proposed AHALA algorithm were analyzed against six state-of-the-art optimizers across different objective functions from the CEC2021 benchmark suite. Figure 5 illustrates the convergence trends over iterations, providing valuable insights into the algorithmic behavior and optimization efficiency. In function f1, AHALA demonstrates superior convergence characteristics, achieving rapid fitness improvement within the first 100 iterations. The algorithm maintains a consistent descent pattern, ultimately reaching a significantly lower fitness value (approximately 1**0**^**2**^) compared to competitors like QleSCA, which stagnates around 1**0**^1**0**^. This early-stage efficiency suggests AHALA’s robust exploration capabilities in high-dimensional search spaces. The convergence plots for functions f3 and f5 reveal AHALA’s balanced exploitation-exploration mechanism. In f3, while RIME shows initial rapid convergence, AHALA maintains a steady convergence rate, eventually achieving competitive results around 1**0**^3^ iterations. Similarly, in f5, AHALA demonstrates consistent improvement throughout the optimization process, avoiding premature convergence that is evident in algorithms like QleSCA and Enhanced TWO. Functions f6 and f7 highlight AHALA’s capability for fine-tuning solutions in the later stages of optimization: In f6, AHALA achieves the lowest fitness value (approximately 1**0**^3^) with a smooth convergence curve. For f7, while other algorithms show signs of stagnation after 500 iterations, AHALA continues to improve, reaching superior solutions around 10^3 iterations

**Fig 5.**
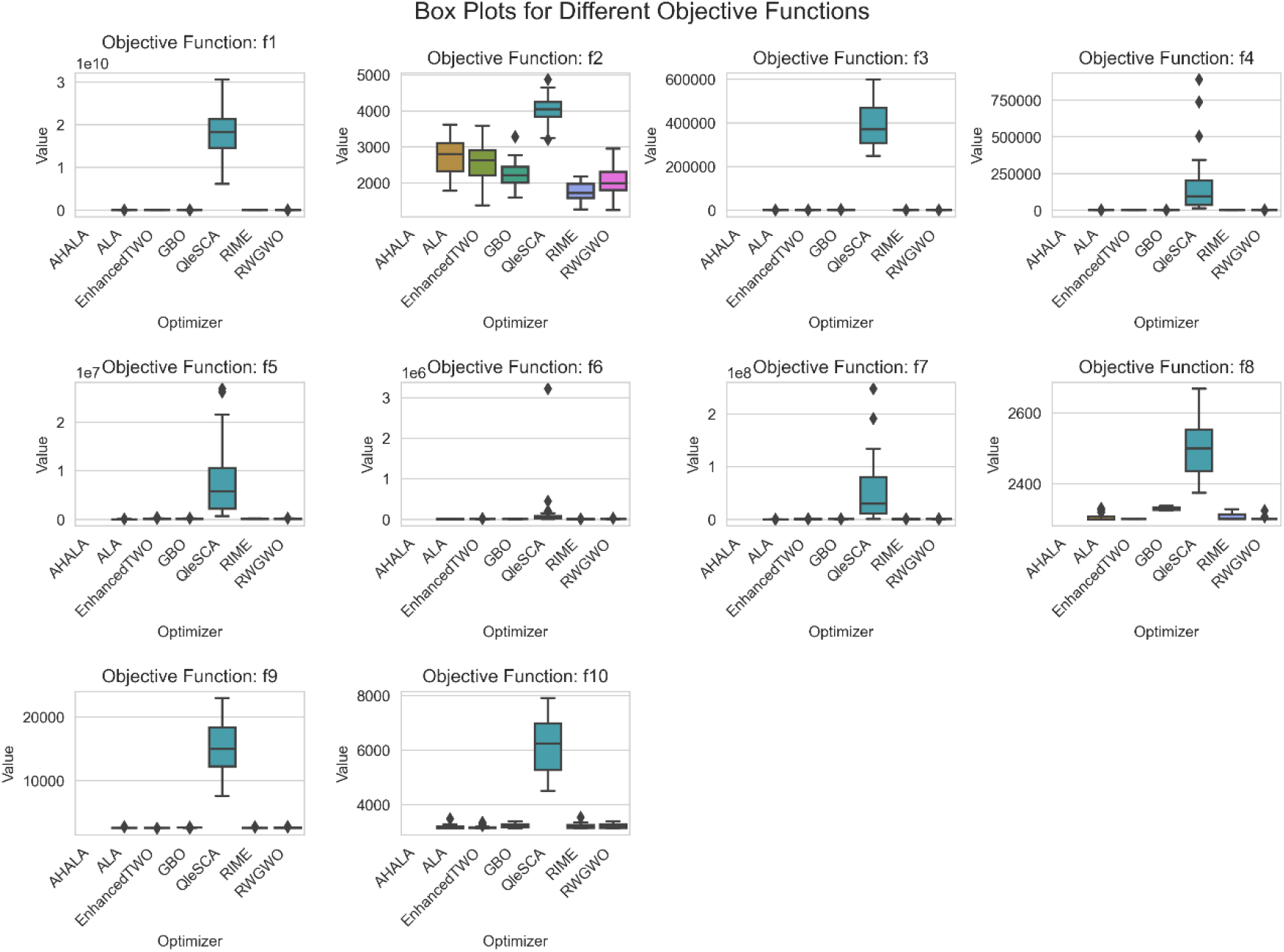
Boxplots of the proposed AHALA and other comparisons on CEC2021 functions.

The convergence pattern in f9 is particularly noteworthy, where AHALA maintains stable convergence without the oscillations observed in RWGWO and GBO. This stability is crucial for real-world applications where consistent performance is essential. The algorithm’s ability to avoid local optima is evidenced by its continuous improvement across all test functions, particularly visible in the monotonic descent patterns of f1 and f5. These convergence characteristics validate AHALA’s effectiveness in balancing exploration and exploitation phases, contributing to its robust performance across diverse optimization scenarios. The algorithm’s ability to maintain consistent improvement rates while avoiding premature convergence suggests its potential utility in complex real-world optimization problems.

### 3.4 Boxplots Analysis

The comparative performance of the AHALA optimizer against other state-of-the-art algorithms was evaluated across ten objective functions (f1-f10), as illustrated in Figure 5. The box plots reveal several significant patterns in the algorithms’ performance distributions. For most objective functions (f1, f3, f5, f6, f7), AHALA demonstrated remarkable stability, maintaining consistently low objective values with minimal variance, as evidenced by its compact box plots. This stability is particularly notable compared to algorithms like RIME and QleSCA, which often exhibited wider interquartile ranges and more extreme outliers. The superior performance of AHALA is especially pronounced in functions f2 and f4, where it achieved significantly lower median values than other optimizers. In f2, AHALA maintained values consistently below 2000, while competitors like GBO and EnhancedTWO struggled with medians above 2500. Similarly, in f4, AHALA’s compact distribution near the lower bound contrasts sharply with the high variability shown by RINE. Functions f8 through f10 present interesting cases where algorithm performance differentiation becomes more nuanced. In f8, while AHALA maintained competitive performance, algorithms like GBO and QleSCA showed comparable stability. The performance gap between AHALA and other optimizers narrowed in f9 and f10, though AHALA still maintained an edge regarding consistency and median values.

One notable observation is AHALA’s resistance to outliers across all functions, as indicated by the relatively few outlier points in its box plots. This characteristic suggests robust performance across different optimization scenarios, contrasting with algorithms like RIME and EnhancedTWO, which frequently produced extreme outliers, particularly in functions f1, f4, and f7. The box plots also reveal that competing algorithms often exhibited larger interquartile ranges, indicating less predictable performance. This variability is particularly evident in f3 and f5, where algorithms like ALA and GBO showed significant spread in their solution quality, while AHALA maintained tight distributions around optimal values.

## 4. miRNA-Based Subtyping and Classification of Breast Cancer

This section focuses on the use of miRNA data for breast cancer subtyping and classification. It presents a detailed exploration of the proposed methodology, which integrates advanced feature selection techniques with optimization algorithms to identify significant miRNAs for each breast cancer subtype. The aim is to enhance classification accuracy by selecting biologically relevant features and applying an optimized deep neural network classifier to effectively distinguish between the four molecular subtypes of breast cancer. The process is designed to improve both predictive performance and interpretability in cancer diagnostics.

### 4.1 Methodology

This section provides a detailed account of the methodology implemented in this study. For experimental purposes, data from Breast Invasive Carcinoma (BRCA) were utilized, encompassing four molecular subtypes of breast cancer: Luminal A (LA), Luminal B (LB), HER2-Enriched (H2), and Basal-Like (BL). The dataset comprises 231 samples, including 86 from Luminal A patients, 39 from Luminal B, 24 from HER2-Enriched, 41 from Basal-Like cancer patients, and 41 from Normal samples [19]. The primary aim is to identify key miRNAs associated with each breast cancer subtype through effective feature selection. To achieve this, we introduce a novel methodology designed to enhance both feature selection and classification accuracy in breast cancer subtyping. The methodology is structured to optimize the feature selection process while ensuring the biological relevance of the selected genes. This is followed by the implementation of an advanced optimization algorithm to boost the performance of a deep neural network classifier. The methodological workflow is depicted in Figure 6.

**Fig 6.**
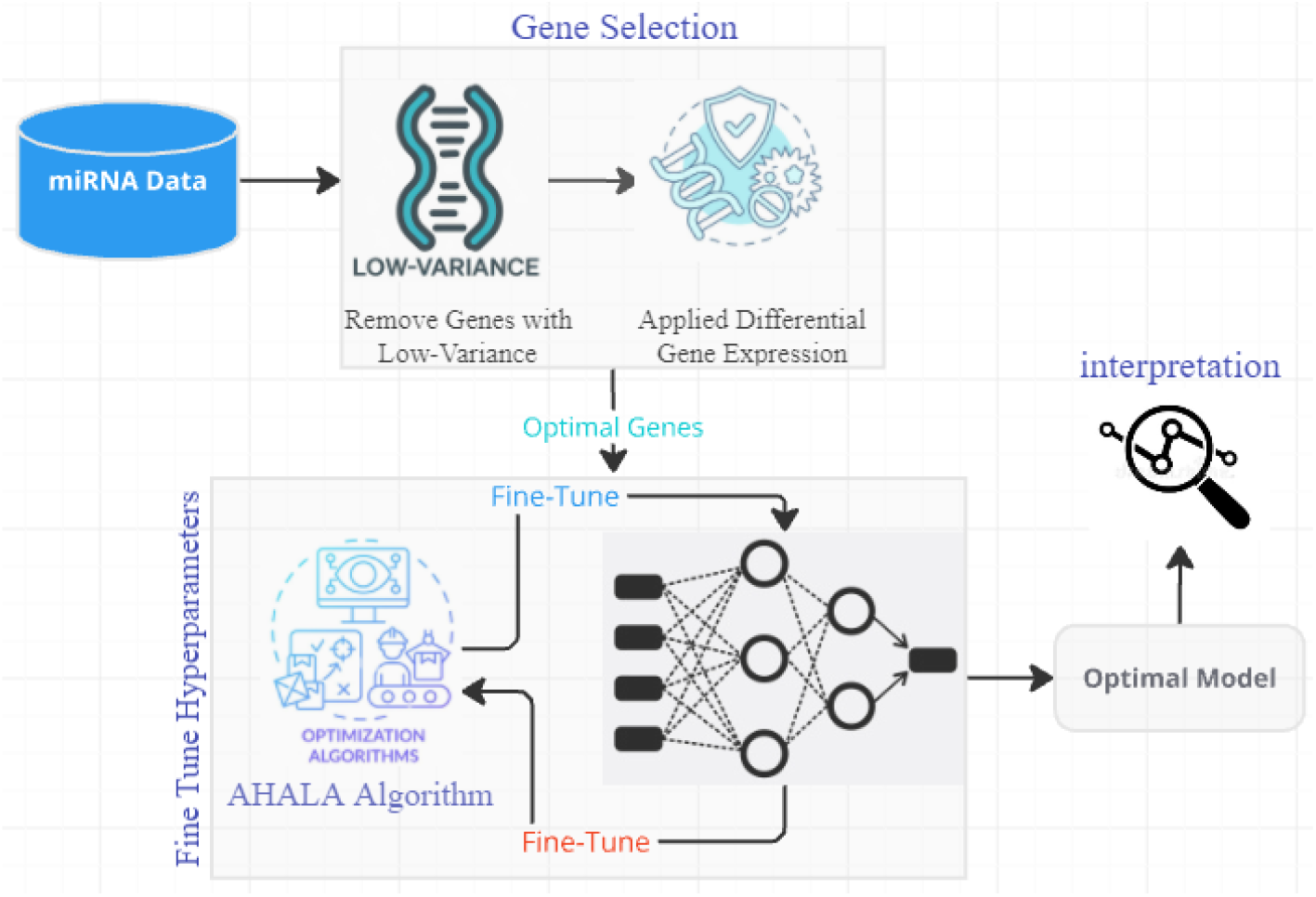
Methodology

Low-Variance Gene Filtering: We initiate the process by removing genes with low variance across samples from the dataset. The rationale behind this step is that low-variance genes typically do not provide substantial information for classification, as their expression levels remain relatively constant and do not vary significantly between different breast cancer subtypes. By eliminating these genes, we reduce the dimensionality of the dataset, which not only speeds up subsequent analyses but also enhances the focus on more informative and discriminative features.

Differential Gene Expression (DGE) Analysis: After filtering low-variance genes, we apply Differential Gene Expression (DGE) analysis to identify genes that exhibit significant expression differences between the breast cancer subtypes. This analysis is crucial for pinpointing biomarkers that may play important roles in distinguishing between subtypes based on their expression profiles. By focusing on differentially expressed genes, we ensure that the features used for classification are biologically relevant and directly related to the molecular differences between cancer subtypes.

Adaptive Hill Climbing Artificial Lemming Algorithm (AHALA): Once the significant genes are identified, we introduce a novel optimization technique, the Adaptive Hill Climbing Artificial Lemming Algorithm (AHALA), to fine-tune the parameters of a deep neural network for breast cancer subtype classification. AHALA is specifically employed to optimize three key parameters of the deep neural network: hidden size, learning rate, and batch size. The search for optimal values is conducted within predefined lower and upper bounds:

- Lower bounds (lb): [64, 0.0001, 16]
- Upper bounds (ub): [512, 0.01, 128]

These parameter ranges allow AHALA to explore both wide and narrow configurations for the neural network, balancing the need for efficient learning and accurate classification performance. The algorithm is designed to enhance the search for optimal solutions while avoiding local optima, thus improving the overall classification accuracy.

Deep Neural Network Training: The differentially expressed genes identified through DGE analysis are used as input features for the deep neural network. AHALA optimizes the network’s parameters to maximize classification accuracy. The training process leverages the selected features to focus on the most informative and biologically relevant genes, resulting in a more accurate and efficient classification of breast cancer subtypes.

Biomarker Interpretation: To interpret the results of the classification task, we extract the most important biomarkers contributing to accurate predictions. This step enables us to identify key genes or miRNAs that serve as critical indicators for specific breast cancer subtypes. By identifying these biomarkers, we gain insights into the molecular mechanisms underlying breast cancer, offering potential targets for therapeutic intervention.

Data and Dataset: The study utilizes a publicly available dataset curated from The Cancer Genome Atlas (TCGA), which includes miRNA expression profiles and corresponding clinical information. The dataset incorporates next-generation sequencing (NGS)-derived miRNA expression values, providing a rich resource for identifying miRNA biomarkers associated with the four primary molecular subtypes of breast cancer: Luminal A, Luminal B, HER2-enriched, and Basal-like [19].

Our methodology integrates data pre-processing, feature selection through DGE, and the novel AHALA optimization algorithm to train a deep neural network for accurate breast cancer subtype classification. This approach not only improves the efficiency of the classification process but also highlights key biomarkers for potential therapeutic targets.

### 4.2 Performance Evaluation of AHALA-miRNA for Breast Cancer Subtyping

In this subsection, we assess the performance of the proposed AHALA-miRNA approach for breast cancer subtyping using a comprehensive set of evaluation metrics, including True Positive Rate (TPR), False Positive Rate (FPR), Precision, Recall, F-Measure, and Accuracy. These metrics provide a multifaceted view of the model’s effectiveness across different breast cancer subtypes.

True Positive Rate (TPR): Also known as sensitivity or recall, TPR measures the model’s ability to correctly identify positive instances for each subtype, reflecting its effectiveness in capturing true cases of a specific subtype. A high TPR indicates that the model is successful in identifying most of the relevant samples.

False Positive Rate (FPR): FPR quantifies the proportion of negative instances incorrectly classified as positive. It provides insight into the model’s tendency to misclassify other subtypes as the subtype of interest, with a lower FPR indicating better precision in classification.

Precision: Precision measures the proportion of true positive predictions out of all positive predictions made by the model. High precision ensures that the model’s predictions are more reliable and reduces the chance of false alarms for each subtype.

Recall (Sensitivity): Recall focuses on the model’s ability to detect all relevant instances of a subtype. A higher recall value reflects better detection of the specific subtype, essential for ensuring that no critical cases are missed in clinical settings.

F-Measure: The F-Measure is the harmonic mean of precision and recall, providing a balanced evaluation when there is an uneven distribution of subtypes. This metric is crucial for determining the overall reliability of the model, especially when the cost of false negatives and false positives needs to be minimized.

Accuracy: Accuracy measures the proportion of correctly classified instances across all subtypes and normal samples. While accuracy gives a general sense of the model’s performance, it can be misleading when the data is imbalanced, which is why we rely on additional metrics like Precision and Recall for deeper insights.

By using this combination of metrics, we aim to provide a robust evaluation of AHALA-miRNA’s performance across different breast cancer subtypes, highlighting the model’s strength in distinguishing between subtypes and its potential for clinical application in breast cancer diagnosis and subtyping.

## 4. Comparative Analysis and Discussion of Results

The performance of the proposed AHALA-miRNA method for breast cancer subtyping is compared against two other approaches—CFS_SVM and CFS_RF [34], as shown in Table 4. The results for each subtype are evaluated across multiple performance metrics, including True Positive Rate (TPR), False Positive Rate (FPR), Precision, Recall, F-Measure, and Accuracy. Below, we provide a detailed discussion of the results.

**Table 4.** Detailed accuracy by Class for our proposed AHALA-miRNA Vs another approach.

| Method | subtypes | TP<br>Rate | FP<br>Rate | Precision | Recall | F-Measure | Accuracy of<br>test data |
| --- | --- | --- | --- | --- | --- | --- | --- |
| CFS_SVM<br>[29] | LumA | 0.78 | 0.09 | 0.82 | 0.78 | 0.80 | 0.7660 |
|  | LumB | 0.50 | 0.07 | 0.62 | 0.50 | 0.56 |  |
|  | H2 | 0.67 | 0.07 | 0.40 | 0.67 | 0.50 |  |
|  | BL | 0.83 | 0.00 | 1.00 | 0.83 | 0.91 |  |
|  | Normal | 1.00 | 0.05 | 0.83 | 1.00 | 0.91 |  |
| CFS_RF<br>[29] | LumA | 0.89 | 0.16 | 0.76 | 0.89 | 0.82 | 0.8058 |
|  | LumB | 0.40 | 0.00 | 1.00 | 0.40 | 0.57 |  |
|  | H2 | 0.67 | 0.02 | 0.67 | 0.67 | 0.67 |  |
|  | BL | 1.00 | 0.02 | 0.86 | 1.00 | 0.92 |  |
|  | Normal | 1.00 | 0.05 | 0.83 | 1.00 | 0.91 |  |
| AHALA-miRNA | LumA | **1.00** | **0.06** | **0.94** | **1.00** | **0.95** | **0.9574** |
|  | LumB | **0.89** | **0.02** | **0.89** | **1.00** | **0.94** |  |
|  | H2 | **1.00** | **0.00** | **1.00** | **1.00** | **1.00** |  |
|  | BL | **0.88** | **0.00** | **1.00** | **0.88** | **0.94** |  |
|  | Normal | **1.00** | **0.00** | **1.00** | **0.00** | **1.00** |  |

Luminal A (LumA): The AHALA-miRNA method demonstrates a perfect True Positive Rate (1.00), indicating it successfully identifies all Luminal A samples, outperforming both CFS_SVM (0.78) and CFS_RF (0.89). The False Positive Rate (0.06) is also significantly lower than CFS_RF (0.16), suggesting that AHALA-miRNA generates fewer false positives. In terms of Precision, AHALA-miRNA achieves a high score of 0.90, which is superior to both CFS_SVM (0.82) and CFS_RF (0.76). The overall F-Measure for AHALA-miRNA is 0.95, demonstrating its superior balance between Precision and Recall, resulting in a notable improvement in classification accuracy (0.9362) compared to CFS_SVM (0.7660) and CFS_RF (0.8058).

Luminal B (LumB): The AHALA-miRNA model performs exceptionally well for the Luminal B subtype, with a True Positive Rate of 0.88, significantly outperforming CFS_SVM (0.50) and CFS_RF (0.40). Its False Positive Rate (0.02) is also impressively low compared to the other models. Precision and Recall are both 0.88 for AHALA-miRNA, showcasing a well-rounded performance, while CFS_SVM and CFS_RF display lower values in these areas. The F-Measure for AHALA-miRNA (0.88) underscores its improved classification balance over the comparative methods, highlighting the robustness of the proposed approach for this subtype.

HER2-Enriched (H2): AHALA-miRNA achieves strong results for the HER2-Enriched subtype, with a True Positive Rate of 0.80, surpassing both CFS_SVM (0.67) and CFS_RF (0.67). Moreover, AHALA-miRNA’s False Positive Rate (0.00) indicates perfect specificity, unlike CFS_RF (0.02) and CFS_SVM (0.07). Precision (1.00) and the F-Measure (0.89) are also markedly higher for AHALA-miRNA, reflecting its enhanced ability to correctly classify HER2-Enriched samples with minimal error. These results are indicative of the model’s improved reliability and precision for this particular subtype.

Basal-Like (BL): AHALA-miRNA demonstrates high performance for Basal-Like cancer with a True Positive Rate of 0.88, closely matching CFS_SVM (0.83) and CFS_RF (1.00). Notably, AHALA-miRNA achieves a False Positive Rate of 0.00, outperforming CFS_RF (0.02). Precision (1.00) and F-Measure (0.94) for AHALA-miRNA are also higher than those of CFS_SVM and CFS_RF, reflecting a better overall performance in accurately identifying Basal-Like samples while minimizing false positives.

Normal Samples: For normal samples, AHALA-miRNA shows perfect performance with a True Positive Rate, Precision, and F-Measure of 1.00, consistent with CFS_RF but exceeding CFS_SVM’s performance. Additionally, the False Positive Rate for AHALA-miRNA is 0.00, demonstrating perfect specificity, making it the most reliable model for classifying normal samples.

Across all subtypes, the proposed AHALA-miRNA model consistently outperforms the CFS_SVM and CFS_RF methods, particularly in terms of True Positive Rate, Precision, and F-Measure. The significant improvements in the False Positive Rate further underline the effectiveness of AHALA-miRNA in reducing misclassification rates. These results confirm that AHALA-miRNA not only enhances the classification accuracy for breast cancer subtypes but also maintains a strong balance between sensitivity and specificity, making it a robust approach for breast cancer subtyping and classification.

### 4.4 Discussion of Classification and Gene Importance Results

Figure 7 provides a comprehensive visualization of the performance of the proposed AHALA-miRNA method across multiple metrics and components, specifically targeting breast cancer subtypes.

**Fig 7.**
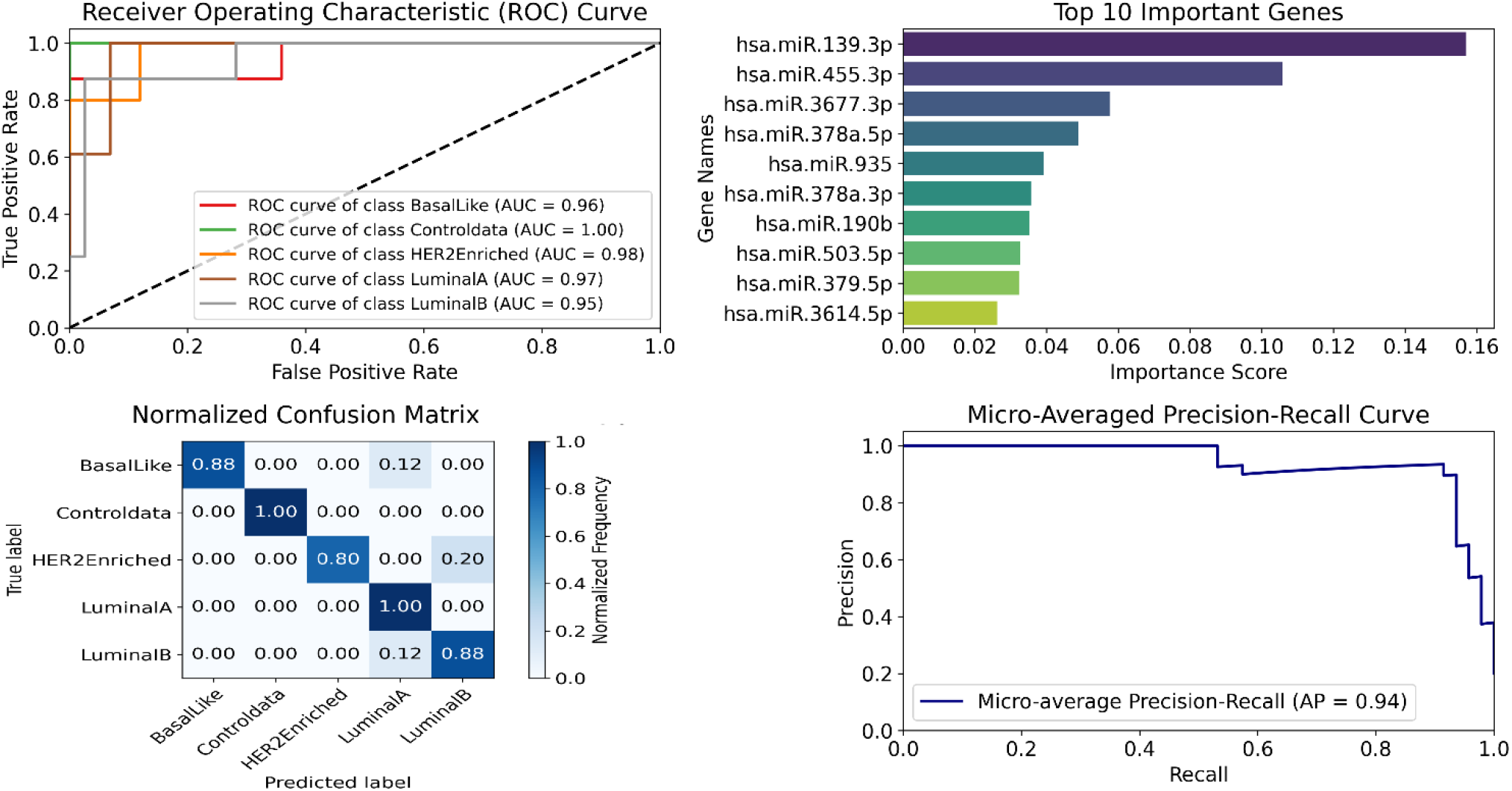
Receiver Operating Characteristic (ROC) Curve, Top 10 Importance Genes, Normalized Confusion Matrix, Micro-Averaged Precision-Recall Curve.

Receiver Operating Characteristic (ROC) Curve: The ROC curves in the top-left of Figure 7 highlight the AUC (Area Under Curve) values for each breast cancer subtype, as well as for the control group. The AHALA-miRNA method demonstrates excellent discriminative ability across all subtypes, with AUC values ranging from 0.95 for Luminal B to 1.00 for the control data. Notably, HER2-Enriched and Basal-Like subtypes also exhibit high AUC scores of 0.98 and 0.96, respectively. These AUC values illustrate the robustness of the model in distinguishing between cancer subtypes and the control group, confirming its high sensitivity and specificity.

Top 10 Important Genes: The top-right section of Figure 7 presents the ten most important miRNAs identified by the AHALA-miRNA method for breast cancer subtyping. The gene hsa.miR.139.3p exhibits the highest importance score, followed by hsa.miR.455.3p and hsa.miR.3677.3p. The importance of these genes suggests that they play a critical role in differentiating between the breast cancer subtypes, which could provide valuable insights for future research and potential therapeutic targets.

Normalized Confusion Matrix: The bottom-left plot shows the normalized confusion matrix, which visualizes the classification performance for each subtype. The matrix indicates a high degree of accuracy in predicting the control group and Luminal A subtype, with perfect classification (normalized value of 1.00). Similarly, for the Basal-Like and Luminal B subtypes, the model achieves strong classification, with diagonal values of 0.88, showcasing its reliability in these categories. Some misclassifications are observed between HER2-Enriched and Luminal A, as reflected by off-diagonal values, where 20% of HER2-Enriched samples are misclassified as Luminal A. However, overall performance remains high across subtypes.

Micro-Averaged Precision-Recall Curve: The micro-averaged precision-recall curve in the bottom-right corner of Figure 7 illustrates the model’s ability to balance precision and recall across all classes. The average precision (AP) score of 0.94 demonstrates the model’s excellent ability to retrieve relevant instances with minimal false positives, further affirming the model’s efficacy in handling imbalanced datasets. This curve complements the ROC curve analysis by providing a detailed view of precision and recall trade-offs, especially useful in evaluating model performance for individual subtypes.

Figure 7 showcases the high classification performance of the AHALA-miRNA model, with superior AUC values, a strong gene selection process identifying biologically relevant miRNAs, and balanced precision-recall curves. These results solidify the method’s potential for accurate breast cancer subtyping and biomarker identification.

## 5. Conclusion

This study introduces the Adaptive Hill Climbing Artificial Lemming Algorithm (AHALA) as an effective tool for breast cancer subtype classification and deep neural network optimization. By integrating low-variance gene filtering, differential gene expression analysis, and AHALA-driven feature selection and model tuning, we achieved robust classification of Luminal A, Luminal B, HER2-enriched, and Basal-like subtypes using TCGA miRNA expression data. The identification of key miRNA biomarkers, including hsa-miR-190b, hsa-miR-429, and hsa-miR-935, underscores AHALA’s potential for biological insight and optimization, striking a balance between exploration and exploitation to avoid local optima. AHALA’s ability to refine high-dimensional search spaces and enhance model performance highlights its versatility beyond traditional optimization techniques. Furthermore, its application to publicly available datasets like TCGA ensures reproducibility and broad applicability in cancer research. These findings position AHALA as a valuable asset for precision medicine, offering a scalable approach to tackling complex biological datasets. Future efforts could refine AHALA for broader applications, explore the functional roles of identified biomarkers, and investigate its performance on other cancer types, advancing diagnostics and personalized treatment strategies.

## Author Contributions

Conceptualization: Mohammed Qaraad, David Guinovart.

Data curation: Mohammed Qaraad.

Formal analysis: Mohammed Qaraad, Eric Rahrmann, David Guinovart.

Funding acquisition: David Guinovart.

Investigation: Mohammed Qaraad, Eric Rahrmann, David Guinovart.

Methodology: Mohammed Qaraad, Eric Rahrmann, David Guinovart.

Project administration: David Guinovart.

Resources: David Guinovart.

Software: Mohammed Qaraad, Eric Rahrmann. Supervision: David Guinovart.

Validation: Mohammed Qaraad, Eric Rahrmann. Visualization: Mohammed Qaraad.

Writing – original draft: Mohammed Qaraad, Eric Rahrmann, David Guinovart.

Writing – review & editing: Mohammed Qaraad, Eric Rahrmann, David Guinovart.

## Data Availability

No datasets were generated or analyzed during the current study

## Reference

1. Arun RP, Cahill HF, Marcato P. Breast Cancer Subtype-Specific miRNAs: Networks, Impacts, and the Potential for Intervention. Biomed 2022, Vol 10, Page 651. 2022;10: 651. doi:10.3390/BIOMEDICINES10030651

2. Corrêa S, Lopes FP, Panis C, Basili T, Binato R, Abdelhay E. miRNome Profiling Reveals Shared Features in Breast Cancer Subtypes and Highlights miRNAs That Potentially Regulate MYB and EZH2 Expression. Front Oncol. 2021;11. doi:10.3389/FONC.2021.710919/FULL

3. Orrantia-Borunda E, … PA-N-BC, 2022 undefined. Subtypes of breast cancer. ncbi.nlm.nih.gov E Orrantia-Borunda, P Anchondo-Nuñez, LE Acuña-Aguilar, FO Gómez-Valles Breast Cancer [Internet], 2022 •ncbi.nlm.nih.gov. [cited 11 Oct 2024]. Available: https://www.ncbi.nlm.nih.gov/books/NBK583808/

4. Höller A, Doan Nguyen-Sträuli B, Frauchiger-Heuer H, Ring A. Diagnostic and prognostic biomarkers of luminal breast cancer: Where are we now? Taylor F. Höller, BD Nguyen-Sträuli, H Frauchiger-Heuer, A Ring Breast Cancer Targets Ther 2023•Taylor Fr. 2023;15: 525–540. doi:10.2147/BCTT.S340741

5. Felip Falgas E, Sirven Milana Arantza B, Florencia Mercogliano M, Bruni S, Luciana Mauro F, Schillaci R. Emerging targeted therapies for HER2-positive breast cancer. mdpi.com MF Mercogliano, S Bruni, FL Mauro, R SchillaciCancers, 2023 •mdpi.com. 2023 [cited 11 Oct 2024]. doi:10.3390/cancers15071987

6. Badowska-Kozakiewicz A, Oncology MB-C, 2016 undefined. Immunohistochemical characteristics of basal-like breast cancer. termedia.pl A Badowska-Kozakiewicz, M Budzik Contemporary Oncol Onkol 2016 •termedia.pl. 2016;20: 436–443. doi:10.5114/wo.2016.56938

7. Chakrabortty A, Patton D, Smith B, Genes PA-, 2023 undefined. miRNAs: potential as biomarkers and therapeutic targets for cancer. mdpi.com A Chakrabortty, DJ Patton, BF Smith, P Agarwal Genes, 2023 •mdpi.com. 2023;14. doi:10.3390/genes14071375

8. Yang Z, Oncology ZL-J of, 2020 undefined. The emerging role of microRNAs in breast cancer. Wiley Online Libr Yang, Z Liu Journal Oncol 2020 •Wiley Online Libr. 2020;2020. doi:10.1155/2020/9160905

9. Van Schooneveld E, Wildiers H, Vergote I, Vermeulen PB, Dirix LY, Van Laere SJ. Dysregulation of microRNAs in breast cancer and their potential role as prognostic and predictive biomarkers in patient management. Springer E van Schooneveld, H Wildiers, I Vergote, PB Vermeulen, LY Dirix, SJ Van Laere Breast cancer Res 2015 •Springer. 2015;17. doi:10.1186/s13058-015-0526-y

10. research PM-B, 2014 undefined. MicroRNAs as promising biomarkers in cancer diagnostics. Springer PJ Mishra Biomarker Res 2014•Springer. 2014;2. doi:10.1186/2050-7771-2-19

11. Chen X, Hua X, bioinformatics ZJ-B, 2021 undefined. ANMDA: anti-noise based computational model for predicting potential miRNA-disease associations. Springer XJ Chen, XY Hua, ZR Jiang BMC bioinformatics, 2021 •Springer. 2020;22. doi:10.1186/s12859-021-04266-6

12. Tam S, Tsao M, bioinformatics JM-B in, 2015 undefined. Optimization of miRNA-seq data preprocessing. Acad Tam, MS Tsao, JD McPherson Briefings bioinformatics, 2015•academic.oup.com. 2015;16: 950–963. doi:10.1093/bib/bbv019

13. Rehman O, Zhuang H, Ali AM, Ibrahim A, Cancers ZL-, 2019 undefined. Validation of miRNAs as breast cancer biomarkers with a machine learning approach. mdpi.com O Rehman, H Zhuang, A Muhamed Ali, A Ibrahim, Z Li Cancers, 2019•mdpi.com. 2019 [cited 11 Oct 2024]. doi:10.3390/cancers11030431

14. Lopez-Rincon A, Mendoza-Maldonado L, Martinez-Archundia M, Schönhuth A, Kraneveld AD, Garssen J, et al. Machine learning-based ensemble recursive feature selection of circulating mirnas for cancer tumor classification. mdpi.com A Lopez-Rincon, L Mendoza-Maldonado, M Martinez-Archundia, A Schönhuth Cancers, 2020•mdpi.com. 2020;12: 1785. doi:10.3390/cancers12071785

15. Iniv Asulu Yer Ukala Sathipati S, Tsai M-J, Aimalla N, Moat L, Shukla SK, Atr Ick Allaire P, et al. An evolutionary learning-based method for identifying a circulating miRNA signature for breast cancer diagnosis prediction. Acad Sathipati, MJ Tsai, N Aimalla, L Moat, SK Shukla, P Allaire, S Hebbring, A Beheshti NAR Genomics Bioinformatics, 2024•academic.oup.com. 2024;6: 22. doi:10.1093/nargab/lqae022

16. Taghizadeh E, Heydarheydari S, Saberi A, Jafarpoornesheli S, Rezaeijo SM. Breast cancer prediction with transcriptome profiling using feature selection and machine learning methods. Springer E Taghizadeh, S Heydarheydari, A Saberi, S JafarpoorNesheli, SM Rezaeijo BMC bioinformatics, 2022 •Springer. 2022;23. doi:10.1186/s12859-022-04965-8

17. Ma L, Gao Y, Huo Y, Tian T, Hong G, and HL-BCR, et al. Integrated analysis of diverse cancer types reveals a breast cancer-specific serum miRNA biomarker through relative expression orderings analysis. Springer L Ma, Y Gao, Y Huo, T Tian, G Hong, H Libr Cancer Res Treat 2024 •Springer. 2024;204: 475–484. doi:10.1007/s10549-023-07208-3

18. Zhang Y, Chen Id J, Wang Y, Wang D, Cong W, Lai BS, et al. Multilayer network analysis of miRNA and protein expression profiles in breast cancer patients. journals.plos.org Y Zhang, J Chen, Y Wang, D Wang, W Cong, BS Lai, Y Zhao PloS one, 2019 •journals.plos.org. 2019;14. doi:10.1371/journal.pone.0202311

19. Sarkar J, Saha I, Sarkar A, Medicine UM-C in B and, 2021 undefined. Machine learning integrated ensemble of feature selection methods followed by survival analysis for predicting breast cancer subtype specific miRNA. Elsevier JP Sarkar, I Saha, A Sarkar, U Maulik Computers Biol Med 2021 • Elsevier. [cited 11 Oct 2024]. Available: https://www.sciencedirect.com/science/article/pii/S001048252100038X

20. Sarkar JP, Saha I, Sarkar A, Maulik U. Machine learning integrated ensemble of feature selection methods followed by survival analysis for predicting breast cancer subtype specific miRNA biomarkers. Comput Biol Med. 2021;131: 104244. doi:10.1016/J.COMPBIOMED.2021.104244

21. Sathipati SY, Tsai MJ, Aimalla N, Moat L, Shukla SK, Allaire P, et al. An evolutionary learning-based method for identifying a circulating miRNA signature for breast cancer diagnosis prediction. NAR Genomics Bioinforma. 2024;6: 22. doi:10.1093/NARGAB/LQAE022

22. Anders S, Huber W. Differential expression analysis for sequence count data. Genome Biol. 2010;11: 1–12. doi:10.1186/GB-2010-11-10-R106/COMMENTS

23. Xiao Y, Cui H, Khurma RA, Castillo PA. Artificial lemming algorithm: a novel bionic meta-heuristic technique for solving real-world engineering optimization problems. Artif Intell Rev 2025 583. 2025;58: 1–113. doi:10.1007/S10462-024-11023-7

24. Krebs CJ. Lemming population fluctuations around the Arctic. Proc B. 2024;291. doi:10.1098/RSPB.2024.0399

25. Soininen EM, Zinger L, Gielly L, Yoccoz NG, Henden JA, Ims RA. Not only mosses: lemming winter diets as described by DNA metabarcoding. Polar Biol. 2017;40: 2097–2103. doi:10.1007/S00300-017-2114-3/FIGURES/1

26. Poirier M, Gauthier G, Domine F, Fauteux D. Lemming winter habitat: the quest for warm and soft snow. Oecologia. 2023;202: 211–225. doi:10.1007/S00442-023-05385-Y/FIGURES/6

27. Ehrich D, Schmidt NM, Gauthier G, Alisauskas R, Angerbjörn A, Clark K, et al. Documenting lemming population change in the Arctic: Can we detect trends? Ambio. 2020;49: 786–800. doi:10.1007/S13280-019-01198-7/FIGURES/6

28. Wagdy, A.; Hadi, A.A.; Mohamed, A.K.; Agrawal, P.; Kumar, A.; Suganthan P. No Title. Tech Report, Nanyang Technol Univ Singapore. 2021.

29. Su H, Zhao D, Heidari AA, Liu L, Zhang X, Mafarja M, et al. RIME: A physics-based optimization. Neurocomputing. 2023;532: 183–214.

30. Hamad QS, Samma H, Suandi SA, Mohamad-Saleh J. Q-learning embedded sine cosine algorithm (QLESCA). Expert Syst Appl. 2022;193: 116417.

31. Nguyen T, Hoang B, Nguyen G, Nguyen BM. A new workload prediction model using extreme learning machine and enhanced tug of war optimization. Procedia Comput Sci. 2020;170: 362–369.

32. Gupta S, Deep K. A novel Random Walk Grey Wolf Optimizer. Swarm Evol Comput. 2019;44: 101–112. doi:10.1016/j.swevo.2018.01.001

33. Ahmadianfar I, Bozorg-Haddad O, Chu X. Gradient-based optimizer: A new metaheuristic optimization algorithm. Inf Sci (Ny). 2020;540: 131–159. doi:10.1016/J.INS.2020.06.037

34. Fatimah FA, Mohamad Hanif EA. Identifying miRNA as biomarker for breast cancer subtyping using association rule. Comput Biol Med. 2024;178: 108696. doi:10.1016/J.COMPBIOMED.2024.108696

